# Reproductive tactics, birth timing and the trade-off between risk avoidance and foraging in an income breeder

**DOI:** 10.1101/2023.03.20.533335

**Authors:** Laura Benoit, Nicolas Morellet, Nadège C. Bonnot, Bruno Cargnelutti, Yannick Chaval, Jean-Michel Gaillard, Anne Loison, Bruno Lourtet, Pascal Marchand, Aurélie Coulon, A.J. Mark Hewison

## Abstract

The behavioural trade-off between foraging and risk avoidance is expected to be particularly acute during gestation and lactation, when the energetic demands of reproduction peak. We investigated how female roe deer, an income breeding ungulate, adjust their management of this trade-off during the birth period in terms of foraging activity and habitat use. We showed that activity levels of reproductive females more than doubled immediately following parturition, when energy demand is highest. Moreover, reproductive females increased their use of open habitat during daytime and ranged closer to roads, but slightly further from refuge woodland, compared to non-reproductive females. However, these post-partum modifications in behaviour were particularly pronounced in late-parturient females who adopted a more risk prone tactic, presumably to compensate for the fitness handicap of their late-born offspring. In income breeders, individuals that give birth late may be forced to trade risk avoidance for resource acquisition during peak allocation to reproduction, likely with significant fitness consequences.

## Introduction

A behavioural trade-off occurs when increasing time allocated to a given activity (e.g. avoiding predation) leads to a decrease in the time that can be allocated to a competing activity (e.g. foraging) (Lima & Dill 1990). Natural selection should favour individuals that balance these competing demands through an optimal decision-making process that ultimately maximises individual fitness (Krebs & Davis 1984). Animals are repeatedly faced with the conflicting demands of acquiring sufficient foraging resources, while satisfying other needs such as avoiding mortality risk, defending a territory, or thermoregulating (Krebs 1980; Sih 1980; Mason et al. 2017). In environments that vary across space and time, habitats that are rich in resources are often associated with a higher risk of predation (Rayor & Uetz 1990; Hernández & Laundré 2005), which may fluctuate at both daily (e.g. night vs. day, Gaynor et al. 2018) and seasonal (e.g. hunting vs. non-hunting season for harvested species, Benhaiem et al. 2008; Adam et al. 2016) scales. As a result, individuals have to choose between feeding on high quality resources, which comes with the cost of greater exposure to predators (i.e. a risk-prone foraging tactic), or sacrificing foraging opportunities to reduce mortality risk (i.e. a risk-averse foraging tactic, see Sih 1980; Verdolin 2006). In this context, decision making can have a decisive impact on individual performance (Caro 2005; Møller & Garamszegi 2012).

When faced with this cost-benefit trade-off between foraging and risk avoidance, the optimal response should differ in relation to individual state (McNamara & Houston 1996), for example, between reproductive and non-reproductive females within a given population (Hamel & Côté 2008; Bjørneraas et al. 2011). Specifically, in mammals, reproductive females must satisfy the pronounced increase in energy demand associated with late gestation and early lactation, while simultaneously avoiding predation of both themselves and their highly vulnerable newborn offspring (Sibly & Brown 2009). In capital breeders (sensu Jönsson 1997), these elevated energy requirements of reproduction are substantially offset by stored energy that is accumulated prior to gestation (e.g. Miller et al. 2006), whereas income breeders (sensu Jönsson 1997) must increase daily resource acquisition (e.g MacWhirter 1991; Dias et al. 2011; Gélin et al. 2013). Consequently, all else being equal, reproductive females should consistently trade other activities in favour of foraging (Neuhaus & Ruckstuhl 2002; Dunbar & Dunbar 1988), particularly for income breeders. However, reproductive females must also prioritise anti-predatory behaviours, for example, increasing vigilance (Barrett *et al*. 2006; Lashley *et al*. 2014, but see Lima 1988) or avoiding high risk habitats (Marchand et al. 2015), especially just after parturition when their offspring are most vulnerable (Clutton-Brock 1991). How reproductive females optimise the balance between the conflicting demands of increased foraging requirements while minimising predation risk is key to understanding variation in maternal performance (Venturelli et al. 2010).

For a given individual state, the optimal response to the trade-off between foraging and risk avoidance depends on the environmental context. For herbivores, phenological cycles of vegetation determine seasonal variation in the nutritional quality of their resources (Chiyo et al. 2005; Chuine 2010; Abbas et al. 2011). A central tenet of evolutionary ecology is that, in seasonal environments, most animals synchronize birth timing to match the annual peak in resource availability (Bronson 1989), ensuring fast growth and high early survival of their offspring. As heavier individuals survive better over the winter than lighter ones (Clutton- Brock et al. 1992; Gaillard et al. 1997), rapid early growth is advantageous, but offspring body mass at winter onset is constrained by birth date (Côté & Festa-Bianchet 2001). Therefore, to compensate the growth handicap of their offspring, females that give birth late in the season should allocate more time to foraging during the lactation period at the expense of higher risk exposure compared to early breeders.

Empirical evidence to date has demonstrated that reproductive and non-reproductive females have contrasting activity budgets during the parturition period (e.g. Ciuti et al. 2006; Grignoli et al. 2007; Hamel & Côté 2008; Bjørneraas et al. 2011). However, to the best of our knowledge, no previous study has evaluated how the occurrence and timing of birth influence the way females resolve the trade-off between foraging activity and risk avoidance in the wild. To fill this knowledge gap, we analysed an exceptional biologging data set comprising a large number of roe deer (*Capreolus capreolus, N=120*) females that were intensively monitored during the parturition season to reliably assess reproductive status and parturition date. Roe deer females are income breeders, generally giving birth to two offspring (Hewison & Gaillard 2001) that are born heavy and grow fast (Gaillard et al. 1993). As a result, allocation to reproduction in this species is among the highest among large herbivores, (i.e. about 26% more than the average large herbivore species, Robbins & Robbins 1979). Thus, roe deer mothers are particularly constrained to increase foraging activity to offset this pronounced increase in energy requirements during late gestation and early lactation (Mauget et al. 1997). Based on metrics of individual activity and habitat use, we therefore expected reproductive females to increase foraging around parturition at the expense of risk avoidance (H1), especially during the day, when human disturbance is highest (Bonnot et al. 2013). More precisely, compared to non-reproductive females, we expected reproductive females to increase activity markedly due to increased foraging needs during early lactation (H1a). As a result, following parturition, we expected them to use open habitats, which are richer (Hewison et al. 2009; Abbas et al. 2011), but perceived as riskier (Bonnot et al. 2013, Panzacchi et al. 2010), more intensively at the expense of proximity to refuge (H1b). Second, while births are highly synchronized in roe deer (80% occur within about 20 days, Gaillard et al. 1993b; Linnell & Andersen 1998), individual variation in parturition date is substantial, repeatable across years for a given female (Plard et al. 2013), and tightly linked to reproductive performance (Plard et al. 2014a, 2015). We thus expected females that gave birth later in the season to make the best of a bad job by trading risk avoidance for resource acquisition (H2), modifying their activity level (H2a) and habitat use (H2b), more markedly than females that gave birth earlier, in an attempt to compensate the growth handicap of their late-born offspring.

## Methods

### Study site

This study was carried out in a 19,000 ha rural region, Vallons et Coteaux de Gascogne (Zone Atelier PyGar), in the south-west of France (43°16′N, 0°53′E). The landscape is hilly and heterogeneous, with a mix of meadows (37% of the total area), crops (cereals, oilseed and fodder crops, 32%), hedgerows (4%), two large forests and numerous small woodland patches (19%) (see Morellet et al. 2011 for more details). Humans are present throughout the study site, living in small villages, farms and isolated houses, and represent the most important risk of mortality for roe deer, mostly through hunting. Drive hunts using dogs occur regularly from September to January (until 2008) or February (since 2009), while stalking takes place from June to September (mainly males). Roe deer density was estimated using a capture-mark-resighting approach to average around 6.0±0.9 individuals / 100 ha in the mixed landscape

### Data collection

From 2002 to 2018, during late fall and winter (from 16 November to 27 March), roe deer were caught using nets, tranquilised with an intramuscular injection of acepromazine, (calmivet 3cc; Montané et al. 2003) and transferred to a wooden retention box to reduce stress and risk of injury. Juveniles (less than one-year-old) were distinguished from adults by the presence of a tri-cuspid third pre-molar milk tooth (Ratcliffe & Mayle 1992). Finally, we fitted a GPS collar (Lotek 3300 GPS or Vectronic GPS PLUS-1C Store On Board and Vertex plus), or a GPS with GSM capability for remote data transmission (Lotek Small WildCell GSM or Vectronic GPS PLUS Mini-1C).

Since 2004, from direct observations of female behaviour equipped with GPS collars, we caught new-borns by hand between late April and early June (see Monestier et al. 2015 for more details). Because females were monitored daily for their appearance and behaviour, and because new-borns can be accurately aged based on their behaviour and morphology (mean age at capture = 5 days; Jullien et al. 1992), we were able to assign parturition dates with an accuracy of 0-5 days (0-2 days in 91% of cases).

All capture and marking procedures were done in accordance with local and European animal welfare laws (prefectural order from the Toulouse Administrative Authority to capture and monitor wild roe deer and agreement no. A31113001 approved by the Departmental Authority of Population Protection).

### Biologging monitoring

GPS collars were scheduled to obtain a location every six hours over the entire year. In addition, for individuals captured since 2009, a location was taken every hour from March to June in juveniles and from mid-March or mid-April to the end of August in adults. We performed differential correction to improve fix accuracy (Adrados et al. 2002). Based on Bjørneraas et al. (2010), we eliminated potentially erroneous data points (0.01% of the total locations) when consecutive locations indicated a step speed that was biologically unfeasible. Most collars hosted an activity sensor that recorded activity level four times per second as the difference in acceleration between two consecutive measurements along the X (forward/ backward) and the Y (sideways) axes (www.lotek.com and www.vectronic-aerospace.com). These measurements were recorded as mean values per 5-minute interval transformed into a standardised measure ranging between 0 and 255. For Lotek 3300 GPS collars, activity data were also measured over 5-minute intervals across the same range, but as a count of vertical X (sideways) and horizontal Y (forward/backward) movements (see Appendix 1 in Supporting Information).

### Defining behavioural metrics to index foraging activity and risk avoidance

To evaluate individual decision making during the period of intensive maternal allocation, we analysed individual activity level and habitat use, which provide reliable proxies of foraging activity and risk avoidance. We focused on an individual-specific time window of 60 days centred on each female’s observed birth date (i.e. 30 days prior to and 30 days following parturition), spanning the last third of gestation and early lactation which are the costliest phases of reproduction (Sadleir 1969, Clutton-Brock et al. 1989). Indeed, although weaning in roe deer occurs in early autumn (Andersen et al. 1998), the first month is the most critical in terms of offspring energy requirements which determine its fate (see Gaillard et al. 2000 for a review). To distinguish the effects of maternal allocation from seasonal fluctuations in other time-dependent drivers such as vegetation phenology, we contrasted activity levels and habitat use profiles of reproductive females with those of philopatric juvenile females (approx. one year old in May) which almost never give birth (Hewison 1996). For these non-reproductive females, we centred the 60-day window on the median parturition date in the population (12^th^ May).

### Indexing foraging activity from metrics of activity level

We expected reproductive females to increase activity following parturition because of the very high resource requirements linked to intensive maternal allocation (H1a & H2a). When large herbivores are active, they spend the vast majority of their time searching for and consuming food (Beier & Mc Cullough 1990; Geist 1963; Georgii 1981). In particular, roe deer spend around half of their active time foraging (Benhaiem *et al*. 2008; Kröschel *et al*. 2017). To index the intensity of foraging activity, we assessed variation in activity level across the 60-day window at both daily and hourly (circadian) scales. We calculated a proxy of activity level as mean VeDBA_activity_ (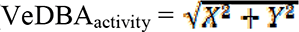, see Benoit et al. 2020) per day and per hour, based on the raw data values recorded over each 5-minute interval by the X and Y activity sensors. This metric has been shown to be a reliable estimator of energy expenditure linked to total body movement (Qasem et al. 2012, Wilson et al. 2020). Because activity values can be influenced by variation in collar fit or sensor type and sensitivity, we scaled the VeDBA_activity_ values for each individual by subtracting the mean of the variable and dividing the result by the variable’s standard deviation (VeDBA_activity_*).

For the above analyses, we retained only those days of monitoring when at least 80% of VeDBA_activity_ records (230 values per day) were available. This analysis was based on a total sample of 72 reproductive females and 40 non-reproductive females (see Appendix 1).

### Indexing foraging activity and risk avoidance from metrics of habitat use

To assess how intensive maternal allocation during gestation and lactation constrained reproductive female habitat use in terms of foraging activity and exposure to risk (H1b & H2b), we computed the proportional use of open habitat, the distance to refuge (woodland) and the distance to roads. As perception of risk may strongly depend on the intensity of human disturbance, which is mostly restricted to daytime (Bonnot et al. 2015), we computed these metrics for daytime only, based on estimated times of sunrise and sunset for each day (sunriset function, maptools package in R software).

First, GPS locations that occurred in habitat types that were rarely used (e.g. roads, gardens, water or orchards, ca. 1% of locations) were removed from the dataset. Then, for each day, we calculated the proportion of all remaining daytime locations that occurred in open habitat (i.e. meadows and crops). As a given threat in exposed situations may be perceived differently depending on proximity to refuge (Padié et al. 2015; Bonnot et al. 2017), we computed the Euclidian distance to the nearest woodland for each daytime location that occurred in open habitat. Finally, to index risk taking behaviour, we computed the Euclidian distance to the nearest road for each daytime location that occurred in open habitat. Distance to the nearest woodland and to the nearest road when in open habitat were weakly correlated (R²: 0.04). This analysis was based on a total sample of 78 reproductive females and 42 non-reproductive females (see Appendix 1).

### Statistical analyses

#### I) Behavioural modifications in relation to reproductive status

We used generalised additive mixed models (GAMMs) with the gamm4 function in the “gamm4” package (Wood & Scheipl 2017) to investigate variation in activity level and habitat use (H1) at the daily and hourly (activity only) scales, as they capture non-linear temporal variations more effectively. To test our hypothesis of state-specific variation in behaviour during the 60-day study window, the most complex model included i/ a separate spline for each reproductive status (reproductive adults vs. non reproductive juveniles) and each period (prior to and following individual date of parturition for reproductive females, or population median date of parturition (12^th^ May) for juvenile females; circadian analysis only); ii/ the additive effects of these factors to control for differences in average behaviour between reproductive and non-reproductive females and between periods. In all models, we included individual date of parturition as an additive effect to control for variation in behaviour linked to phenological changes in the vegetation relative to the timing of maternal care (see Fig S2.1 in Appendix 2) and individual identity as a random effect on the intercept.

##### A) Variation in activity level between reproductive and non-reproductive females

To investigate variation in VeDBA_activity_* at the daily scale, the GAMM included a thin plate regression spline to smooth the effect of time (in days) over the 60-day window. For model selection, we compared a constant model, a model containing only the spline of the day, a model including separate splines of the day for non-reproductive juveniles and reproductive adults, and the three corresponding models that included an additional additive effect of reproductive status.

To investigate variation in VeDBA_activity_* at the hourly scale, we used cyclic cubic splines to smooth the effect of time (in hours) over the 24-hour cycle. For model selection, we compared a constant model, a model containing only the spline of the hour of the day, a model including separate splines of the hour of the day for non-reproductive juveniles and reproductive adults, a model including separate splines for the hour of the day for the two periods, a model including both period-specific and reproductive status-specific temporal effects, and all possible combinations of these models, with or without an additional interaction between these two effects.

##### A) B) Variation in habitat use between reproductive and non-reproductive females

At the daily scale only, we used three sets of GAMMs to investigate variation in the proportional use of open habitat during the day (with a binomial distribution and a logit link function), and in distance to the nearest woodland and the nearest road (square root transformed) during daytime when an individual was located in open habitat. We used a thin plate regression spline to smooth the effect of time (in days) over the 60-day window and the same model set as for analysis of activity level at the daily scale, above.

#### **II)** Behavioural modifications associated with parturition timing

At the daily scale and for reproductive females only, we investigated how modifications in activity level and habitat use varied during each period (prior to and following parturition), in relation to the timing of parturition (H2). In these analyses, we used i/ linear mixed models (package “lme4”, Bates et al. 2015) to investigate variation in daily VeDBA_activity_*, distance to the nearest woodland and distance to the nearest road during daytime when in open habitat; and ii/ generalised linear mixed models (package “lme4”) to investigate proportional daytime use of open habitat (with a binomial distribution and a logit link function). The most complex model included the two-way interaction of parturition date and period. All models included individual identity as a random effect on the intercept.

All models were fitted using maximum likelihood and we used Akaike’s information criterion (AIC, Burnham & Anderson 2002) and Akaike weights to select the model with the strongest support. When ΔAIC between competing models was < 2, following rules of parsimony, we selected and interpreted the model with the fewest parameters. Analyses were performed in R software version 4.0.3 (R Development Core Team, 2020).

## Results

Median parturition date is the 12^th^ May in our study population, but was 16^th^ May [20^th^ April – 11^th^ June] in the sub-sample of GPS monitored females analysed below (see Figure S2.2 in Appendix 2 for the distribution of parturition dates).

### I) Behavioural changes in relation to reproductive status

#### A) Differences in activity level between reproductive and non-reproductive females

The selected model that best described variation in daily activity level included separate smoothed effects of the day for reproductive and non-reproductive females, with an additive effect of reproductive status (see Table S3.1 in Appendix 3). There was a marked difference between reproductive and non-reproductive females in temporal variation of activity level over the 60-day window. In particular, daily VeDBA_activity_* of juveniles decreased over the study period (average Δ daily VeDBA_activity_* ± SE between 30-day periods prior to vs following 12^th^ May of -0.04±0.02). In contrast, in support of H1a, reproductive females markedly increased their activity levels following parturition compared to prior to parturition (average Δ daily VeDBA_activity_* between 30-day periods prior to vs. following parturition of 0.09±0.02, Fig. 1).

**Fig. 1:**
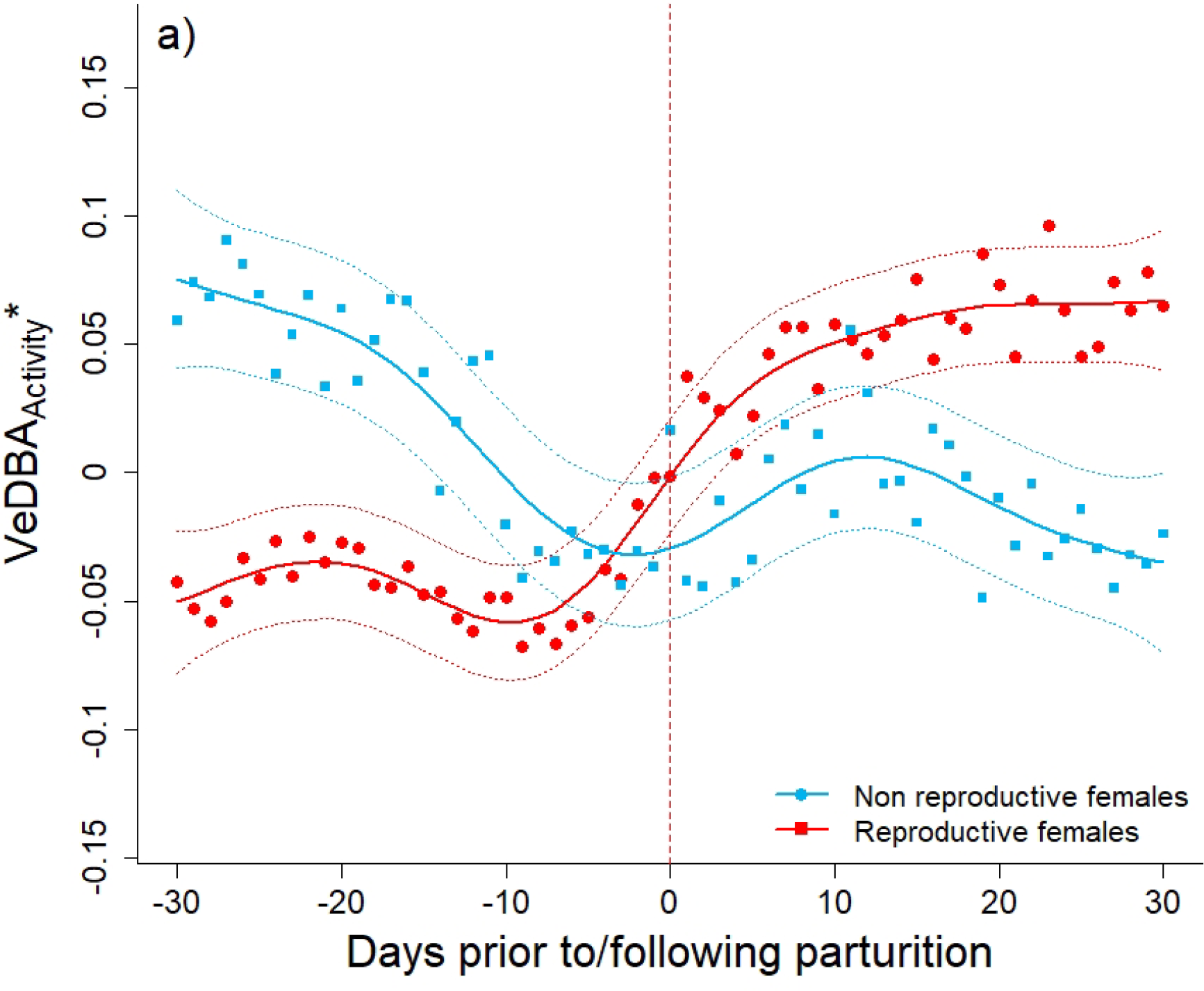
Variation in daily VeDBA_activity_* for the 60-day window centred on individual parturition date for reproductive females (in red) and on the population-level median parturition date (12^th^ May) for non-reproductive juveniles (in blue) Lines (and dotted lines) represent predictions (and their associated 95% confidence intervals) of the selected model (Appendix 3, Table S3.1). Points represent observed values averaged per day for each reproductive status. The vertical dotted line represents the day of parturition.

The selected model that best described the circadian rhythm in activity level contained a separate smoothed effect of the hour of the day for each reproductive status and each period, together with the two-way interaction between period and reproductive status (see Table S3.2 in Appendix 3). Prior to parturition, all females had a similar circadian activity rhythm, with a higher hourly VeDBA_activity_* around dawn and dusk (Fig. 2). These peaks were particularly marked for non-reproductive females. However, following parturition, the observed circadian rhythms differed markedly in relation to reproductive status. First, non-reproductive juveniles maintained a similar overall activity profile across the entire 60-day window (e.g. at 12:00, Δ hourly VeDBA_activity_* prior to vs. following 12^th^ May: 0.01±0.03), although the average level of activity during night-time was slightly lower during the period following 12^th^ May compared to the period prior to this date (e.g. at 00:00, Δ hourly VeDBA_activity_*: -0.14±0.03). In contrast, compared to the pre-parturition period, reproductive females exhibited a substantially higher level of activity following parturition which was especially marked during daytime (Δ hourly VeDBA_activity_* at 12:00: 0.21±0.02) compared to night time (Δ hourly VeDBA_activity_* at 00:00: 0.06±0.02) (H1a).

**Fig. 2:**
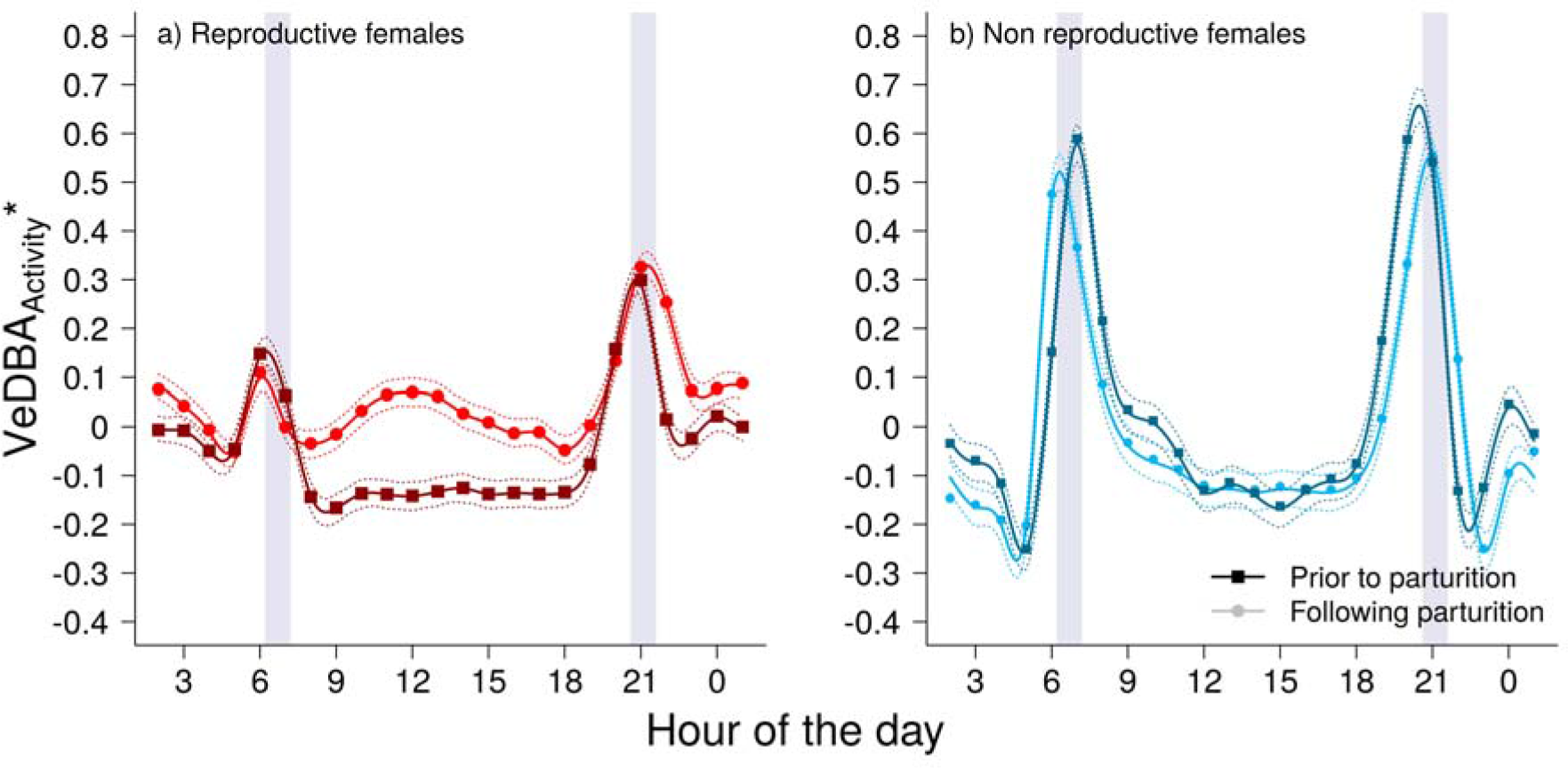
Circadian variation in hourly VeDBA_activity_* for reproductive females (a; in red), and non-reproductive juveniles (b; in blue), prior to (dark colours) and following (light colours) parturition (i.e. individual parturition date for reproductive females, population-level median parturition date (12^th^ May) for non-reproductive juveniles). Full lines represent predictions (and dotted lines their associated 95% confidence intervals) for the selected model (Appendix 3, Table S3.2). Points represent observed values averaged per hour for each period and reproductive status. Grey bars indicate 1 hour around mean twilight over the 60-day window. Hours of the day are in local time (GMT+2).

#### B) Differences in habitat use between reproductive and non-reproductive females

The selected models that best described variation in daytime habitat use over the 60-day window were similar for all three metrics, including a separate smoothed effect of day for each reproductive status, with an additive effect of reproductive status (see Tables S3.3, 3.4 & 3.5 in Appendix 3). First, although the use of open habitat during daytime increased across the 60-day window for all individuals, on average, reproductive females used open habitat more than non-reproductive juveniles (Fig. 3a). In particular, reproductive females were, on average, 1.4 times more often in open habitat during daytime prior to parturition (average proportion of open habitat used: 45±4 % vs. 31±4 % for reproductive and non-reproductive females, respectively), but only 1.1 times more often following parturition (65±4 % vs. 57±5 %) (H1b). Second, reproductive females were, on average, 1.2 times closer to roads during daytime than non-reproductive juveniles, although this difference was slightly more marked following parturition (average distance to the nearest road in open habitat prior to parturition: 216±11 m vs. 271±17 m; following parturition: 205±11 m vs. 271±17 m) (H1b) (Fig. 3b). Finally, all individuals ventured progressively further from woodland during daytime across the 60-day window. However, although reproductive females tended to range, on average, slightly further from woodland than non-reproductive females (average distance to the nearest woodland in open habitat prior to parturition: 67±6 m vs. 58±8 m; following parturition: 81±7 m vs. 78±9 m), there was no consistent difference in relation to reproductive status (Fig. 3c).

**Fig. 3:**
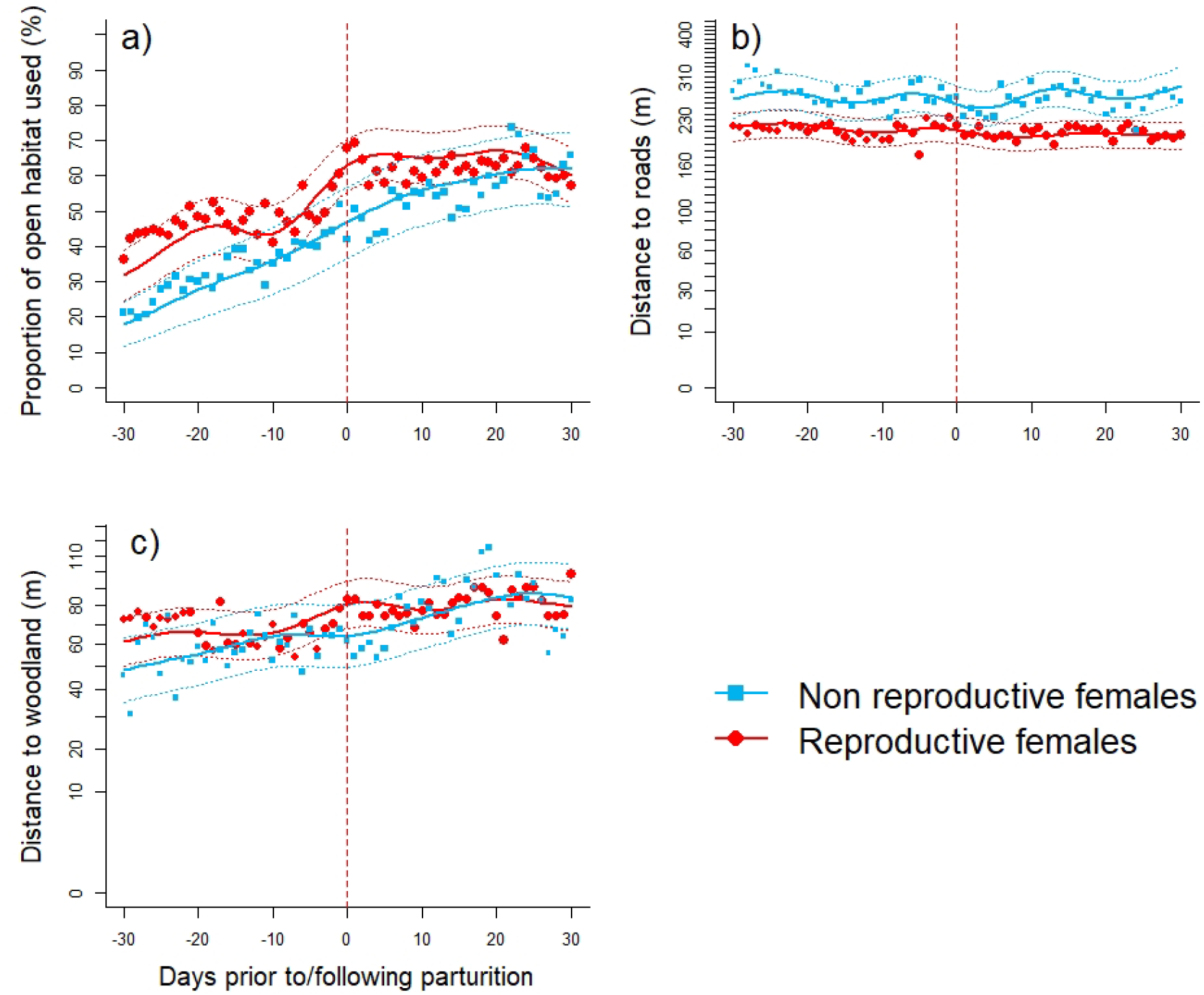
Variation in daytime habitat use across the 60-day window centred on individual parturition date for reproductive females (in red) and on the population-level median parturition date (12^th^ May) for non-reproductive juveniles (in blue), measured as the (a) proportion of open habitat used, (b) distance to the nearest road when in open habitat, and (c) distance to the nearest woodland when in open habitat. Lines (and dotted lines) represent predictions (and their associated 95% confidence intervals) for the selected models (Appendix 3, Tables S3.3, S3.4, S3.5). Points represent observed values averaged per day for each reproductive status. The vertical dotted line represents parturition date.

### II) Behavioural changes in relation to the timing of parturition

#### A) Variation in activity levels of reproductive females in relation to parturition timing

The selected model that best described variation in activity level in relation to parturition date included the two-way interaction between parturition date and period (i.e. prior to vs. following parturition) (Table S4.1 in Appendix S4). Reproductive females that gave birth after the median date of parturition (12^th^ May; “late-parturient females” hereafter) were, on average, less active than females that gave birth prior to this date (“early-parturient females” hereafter). This difference was pronounced prior to parturition (daily VeDBA_activity_* = - 0.08±0.01 for late-parturient females vs. -0.01±0.02 for early-parturient females), but vanished following parturition (0.04±0.01 vs. 0.05±0.02). Hence, as expected (H2a), the individual-level increase in average activity level following parturition was more pronounced for late-parturient females than for early-parturient females. Specifically, late-parturient females increased their average daily VeDBA_activity_* by twice as much as early-parturient females (average Δ VeDBA_activity_* prior to vs. following parturition: 0.12±0.02 vs. 0.06±0.02) (Fig. 4a).

**Fig. 4:**
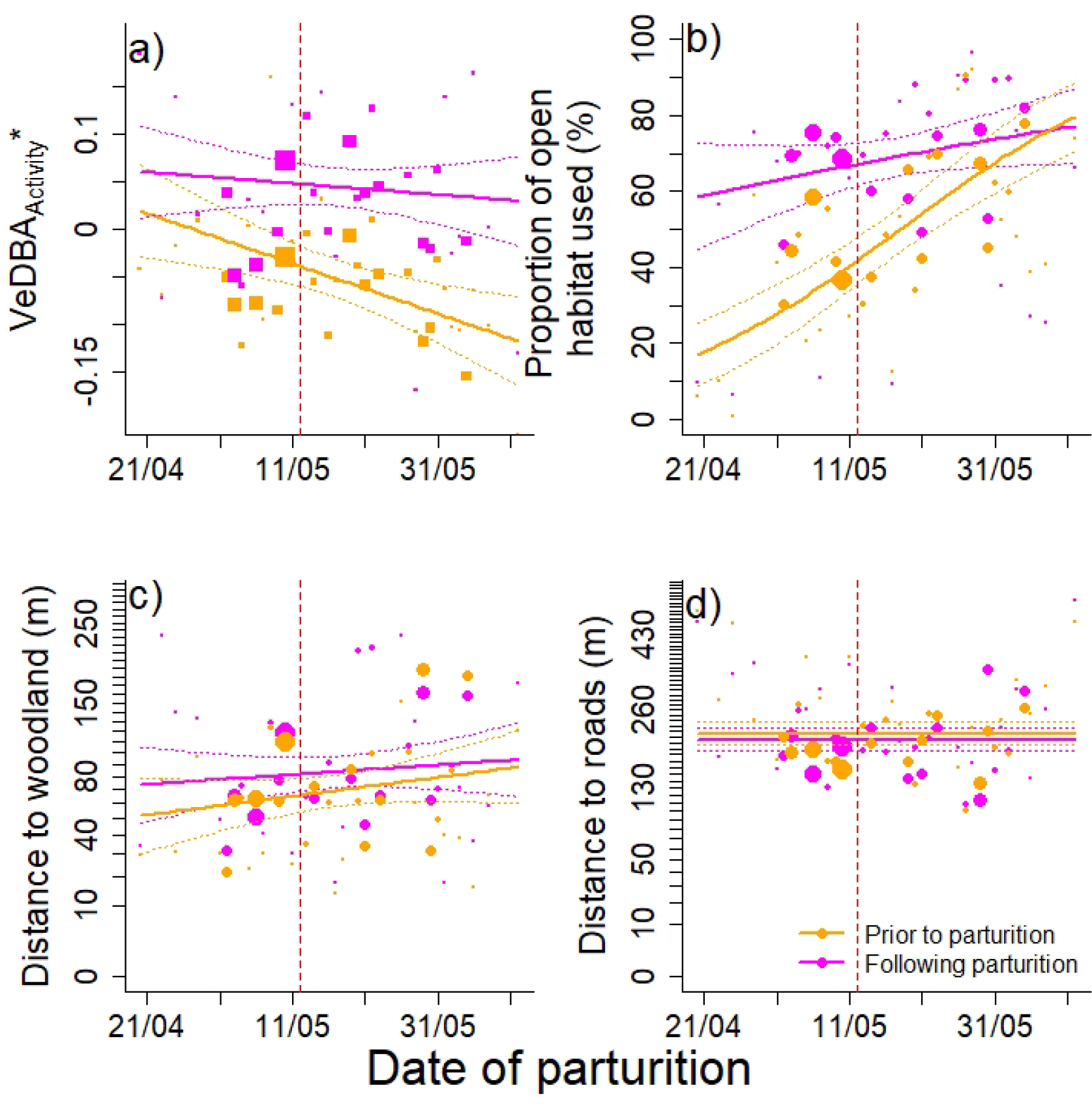
Variation in activity level and daytime habitat use of reproductive females prior to (orange line) and following (purple line) parturition in relation to the timing of parturition: (a) daily VeDBA_activity_*, (b) proportion of open habitat used, (c) distance to the nearest road when in open habitat and (d) distance to the nearest woodland when in open habitat. Lines (and dotted lines) represent predictions (and their associated 95% confidence intervals) for the selected models (Appendix 4, Tables S4.1-4.4). Points represent observed values averaged per period and for females that gave birth on a given day. Point size is proportional to sample size. The vertical dotted line represents the median parturition date in the population (12^th^ May).

#### B) Variation in daytime habitat use of reproductive females in relation to parturition timing

The selected models that best described variation during daytime in i/ the use of open habitat and ii/ the distance to the nearest woodland when in open habitat in relation to parturition date both included the two-way interaction between parturition date and period (prior to vs. following parturition). In contrast, the equivalent model for the distance to the nearest road when in open habitat included the effect of period only (see Table S4.2, 4.3 & 4.4 in Appendix 4). On average, late-parturient females used open habitat during the day more often than early-parturient females (H2b), but this difference was more pronounced prior to parturition (62±4% for late-parturient females vs. 28±4% for early-parturient females) than following parturition (72±3% vs. 63±5%) (Fig. 4b). Thus, following parturition, late- parturient females increased their daytime use of open habitat much less (by 1.2 times) than early-parturient females (by 2.2 times). Similarly, when in open habitat during daytime, late- parturient females ranged, on average, further from woodland than early-parturient females, but more so prior to parturition (77±9 m vs. 58±9 m) than following parturition (88±10 m vs. 77±10 m, Fig. 4c). Finally, parturition date had no effect on the distance to the nearest road when in open habitat during daytime. Irrespective of their parturition date, females tended to range slightly closer to roads during daytime following parturition compared to prior to parturition, but this difference was not marked (average Δ for distance to the nearest road prior to vs. following parturition: -12±15 m, Fig.4d).

## Discussion

For income breeding mammals, the cost-benefit trade-off between foraging and risk avoidance is predicted to be particularly acute during late gestation and early lactation, when the energetic demands of reproduction peak (Lima & Dill 1990; Sadleir 1987). As early survival of offspring is a key component of reproductive success in large herbivores (Gaillard et al. 2000), thereby influencing female fitness, selection should favour a clear-cut behavioural optimum for this trade-off (Venturelli et al. 2010). Using an exceptional biologging data set and intensive field monitoring of reproductive females to establish parturition date, we demonstrated dramatic changes in foraging and risk avoidance behaviour during early lactation in the income breeding roe deer. Furthermore, females exhibited contrasting tactics for balancing risk avoidance versus foraging activity in relation to breeding phenology. Specifically, compared to early breeders, late-breeding females increased post- parturition allocation to foraging activity more markedly, likely in an attempt to compensate the growth handicap of their late-born fawns. Consequently, late-parturient females were forced to adopt a risk prone space use tactic during the energetically critical times of late gestation and early lactation.

### Intensive maternal care in an income breeder drives risk prone behaviour

To offset the energetic costs of peak allocation to reproduction (Oftedal 1985; Gittleman & Thompson 1988), income breeders (sensu Jönsson 1997), such as roe deer, must spend most of their active time searching for and consuming food (Benhaiem *et al*. 2008; Kröschel *et al*. 2017). Here, we obtained strong evidence that, compared to non-reproductive juveniles, reproducing females markedly increased the intensity of their activity immediately following parturition (Fig. 1). This overall response was associated with a marked modification of the circadian rhythm, which is modulated by both physiological state (Daan et al. 2013) and environmental conditions (e.g Pohl 1998). In particular, the increase in activity during early lactation was particularly marked during daytime, when perceived risk is highest (Bonnot et al. 2013), whereas the typical dawn and dusk peaks in activity (Krop-Benesch et al. 2013) were somewhat attenuated (Fig. 2). The tight synchronisation of this dramatic increase in activity with the parturition event of individual females suggests that, for an income breeder, the first weeks of lactation constitute by far the costliest phase of maternal care (see also Clutton-Brock et al. 1989). Intense foraging activity during this period is crucial to sustain rapid offspring growth and high survival, with crucial consequences for fitness (Gaillard et al. 1998; ). Our results indicate that to meet this dramatic increase in energetic requirements, roe deer mothers must remain active throughout the day, alternating between feeding intensively in the richest resource patches within the heterogeneous landscape matrix (Hewison et al. 2009) and providing frequent maternal care for their offspring.

Open habitats are often considered as providing forage of higher quality and quantity for large herbivores during the growing season (Kamler & Homolka 2016), but are often also perceived as riskier (Ciuti et al. 2006; Grignoli et al. 2007; Bjørneraas et al. 2011). Previous studies demonstrated that the use of open habitat increases reproductive success in female roe deer (McLoughlin et al. 2007; Bonnot et al. 2018), although the strength of the response depends on predation pressure (Kjellander et al. 2004) and personality (Monestier et al. 2015; Bonnot et al. 2018). Here, we showed that reproductive females used open habitat during daytime more frequently than non-reproductive females across the entire 60-day study window (Fig. 3a). Thus, around parturition, roe deer mothers prioritise access to rich feeding areas throughout the day, spending up to two thirds of their time in the open, despite the higher associated risk (Jarnemo 2002; Panzacchi et al. 2010). In addition, when feeding in open habitat, reproductive females adopted a risk-prone behaviour, ranging closer to roads (Fig. 3b) and slightly further from refuge woodland (Fig.3c) compared to non-reproductive females. Note that, for both reproductive and non-reproductive females, both the use of open habitat and the distance to woodland increased gradually through the season, from the pre-parturition to the post-parturition phase (Fig. 3). Although juveniles should maximise body growth during this period (Hewison *et al*. 2011), we suggest that the increasing propensity for non-reproductive females to use open habitat and their gradual decrease in activity level (see also Appendix 2) were likely linked to plant phenology. Indeed, plant growth over the course of the spring makes open habitat increasingly attractive for foraging (Abbas et al. 2011), but also less risky once crops and meadows are high enough to provide some degree of cover.

Taken together, our results suggest that the behaviour of roe deer mothers during late gestation and early lactation is primarily driven by the landscape of resources rather than the landscape of fear (Laundré et al. 2001). This contrasts with studies performed on other ungulates where reproductive females prioritise risk avoidance, using poorer but safer habitats (fallow deer: Ciuti et al. 2006; Alpine ibex: Grignoli et al. 2007; moose: Bjørneraas et al. 2011). This apparent discrepancy is likely due to the fact that these species are all much closer to the capital end of the income-capital breeding continuum than roe deer. The lack of significant fat reserves in income breeders, such as roe deer, imposes a strong constraint on females to continually acquire high quality resources to offset the costs of late gestation and early lactation. Failing to do so consistently results in annual reproductive failure, even in the absence of predation (Gaillard et al. 1997). Given these severe constraints, selection should favour risk-prone decision-making tactics. Risk taking is, however, not solely determined by space use (Lima & Dill 1990; Marchand et al. 2014). When avoiding risky areas is not a suitable alternative, females must accommodate increased risk exposure by adjusting their anti-predator behavior, particularly when nursing offspring. Specifically, roe deer females have been reported to remain close to their offspring (Panzacchi et al. 2010), actively defending them (Jarnemo et al. 2004), and potentially to increase their vigilance. Future work should aim to elucidate individual differences in maternal allocation tactics, both in terms of ranging and anti-predator behaviours, and quantify their impact on offspring survival.

### Late-parturient females are more risk-prone than early-parturient females

In mammals, births are scheduled to match closely the seasonal peak in resource availability (Bronson 1989), maximising early survival and growth of offspring. Rapid early growth ensures that juveniles achieve the threshold body size at the onset of winter that is key for subsequent survival prospects (Clutton-Brock et al. 1992; Gaillard et al. 1997). We thus expected females that gave birth late in the season to allocate more intensively during early lactation to compensate for the bad start of their offspring. Our analyses demonstrated that, prior to parturition, average activity level decreased (Fig.4a) and the use of open habitat increased (Fig.4b) with increasingly late parturition date. As a result, pregnant late-parturient females were, on average, less active, but were more frequently in open habitat and further from refuge cover, compared to pregnant early-parturient females, likely linked to vegetation phenology. However, in support of our hypothesis, the increase in allocation to foraging activity immediately following parturition was much more pronounced for late-parturient females than for early-parturient females (H2a). Furthermore, although early-parturient females modified their habitat use following parturition, late-parturient females consistently took more risks by spending more time in open habitat and ranging further from woodland cover. These results support our hypothesis that late-parturient females should seek to compensate the growth handicap of their fawns by dramatically increasing their allocation to foraging activity post-partum, targeting rich open habitat by adopting a risk-prone space use tactic.

Within a given cohort, early-born fawns benefit from better environmental conditions, generating higher survival and performance during adulthood (Plard et al. 2015) through their greater ability to exploit silver spoon effects (sensu Grafen 1988). Hence, the risk-prone behaviour of late-parturient females can be interpreted as a “best of a bad job” tactic to at least partially compensate for the fitness handicap inflicted by their offspring’s bad start. However, in roe deer at least, the decision making of these females appears ultimately to have limited influence on the evolutionary outcome, as late-parturient females have been reported to be of poorer phenotypic quality (Plard et al. 2014b) and to live shorter lives than early-parturient females. Although the spring flush and peak forage productivity is occurring increasingly early over time due to global warming (Parmesan & Yohe 2003), unlike other ungulates (e.g. Pyrenean chamois : Kourkgy *et al*. 2016, reindeer: Paoli *et al*. 2018, red deer: Bonnet *et al*. 2019), roe deer seem unable to adjust their birth timing to match vegetation phenology (Plard et al. 2014a),. Indeed, parturition date is remarkably constant and repeatable over the lifetime of a given female in this species (Plard et al. 2013). As a result, females that consistently give birth late throughout their lifetime will likely pay even higher costs in the future due to this increasingly pronounced phenological mismatch, with obvious negative impacts for their fitness (Visser & Gienapp 2019).

## Acknowledgement

We would like to thank the local hunting associations with the Fédération Départementale des Chasseurs de la Haute Garonne, as well as numerous co-workers and volunteers for their assistance. This project was supported by ‘MovLIt’ Agence Nationale de la Recherche grant ANRL16LCE02L0010L02 to A.J.M.H., A.L. and J.M.G.

