## Appendix for "Reproductive tactics, birth timing and the trade-off between risk avoidance and foraging in an income breeder"

**Appendix S1: GPS and activity data characteristics.**

**Table S1:** Characteristics for GPS and activity data. Numbers are sample sizes (number of females).

|  | GPS data | Activity data | |
| --- | --- | --- | --- |
| Sampling frequency | 6 hours or 1hour | 5 min | |
|  |  | X (sideways) and Y (forward/backward) | Y (sideways) and X (forward/backward) |
| Lotek Small WildCell | 53 | / | 50 |
| Lotek 3300 GPS | 67 | 62 | / |
| Total non-reproductive females | 42 | 13 | 27 |
|  |  | 40 | |
| Total reproductive females | 78 | 49 | 23 |
|  |  | 72 | |
| Total females | 120 | 62 | 50 |
|  |  | 112 | |

**Appendix S2: On the need to account for parturition date in the analyses of behavioural modifications during the period of intensive maternal allocation**

At the population level, average activity level tended to decrease over the period of intensive maternal allocation at both daily (slope: -0.002 for daily VeDBA_activity_*, see Figure S2), and hourly (slope: -0.002 for hourly VeDBA_activity_*) scales. However, both the proportion of open habitat used during the day and the distance to the nearest woodland when in open habitat tended to increase over the same period (odds ratio: 0.035 for proportion of open habitats used; slope: 0.030 for square root transformed distance to the nearest woodland). Finally, distance to the nearest road when in open habitat did not change during this period (slope: -3.247e-05). Thus, since parturition date influences the behavioural measurements (activity level and habitat use), we set the date of parturition to May 12 (median date of parturition for the population) to study behavioural modifications in relation to reproductive status during the 60-day window at daily or hourly (for VeDBA activity only) scales.

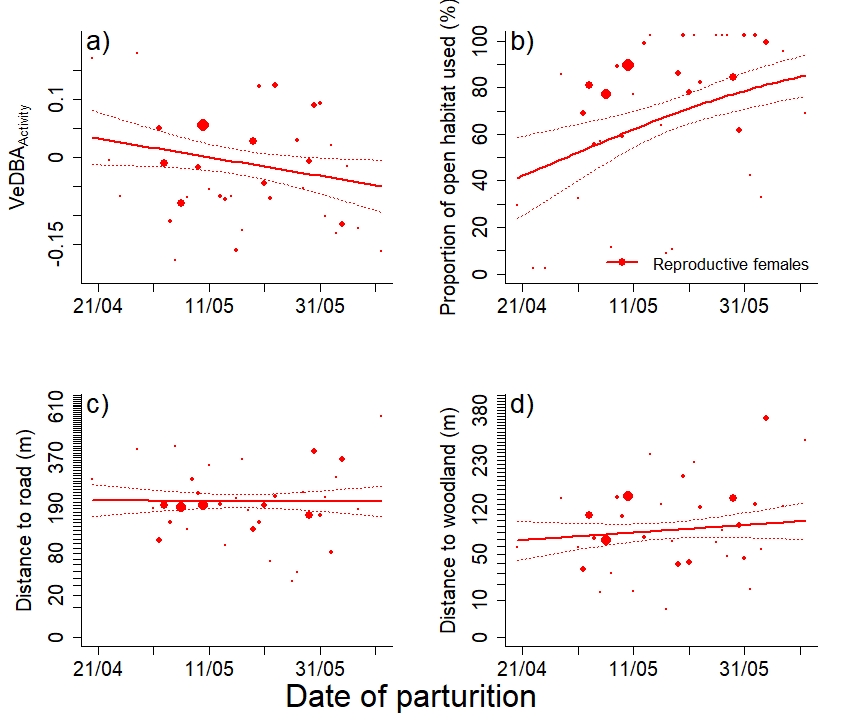
 **Figure S2.1:** Variation in (a) daily VeDBA_activity_, (b) proportion of open habitat used during daytime, (d) distance to the nearest road when in open habitat during daytime and (e) distance to the nearest woodland when in open habitat during daytime, as a function of parturition date of females, considering the other parameters fixed with the variable day set to the observed parturition date and status set to reproductive females. Point size is proportional to sample size. Points (and bars) represent predictions (and associated 95% confidence intervals) of the retained models.

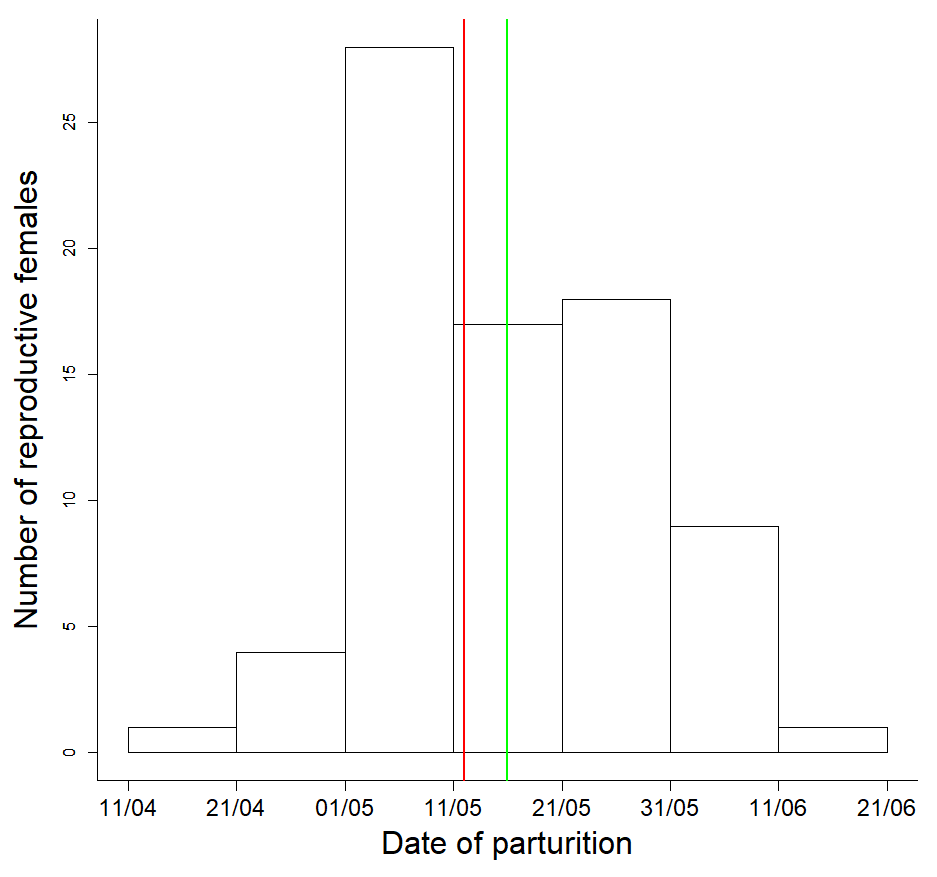
**Figure S2.2:** Distribution of parturition date in the sub-sample of GPS monitored females of our population. The median parturition date of the population (12^th^ May) is represented by a red vertical line and the median parturition date of the sub-sample of GPS monitored females (16^th^ May) is represented by a green vertical line.

**Appendix S3: Model selection with AIC criteria for the analysis of behavioural modifications linked to reproductive status**

Models were ranked according to AIC selection criteria using the difference in the values of AIC (∆AIC), the number of estimated parameters (K) and Akaike weights. The retained model is in bold.

**Table S3.1:** Support for the candidate generalised additive mixed models explaining variation in daily VeDBA_activity_*.

| **Models** | **K** | **AIC** | **∆AIC** |
| --- | --- | --- | --- |
| **s(Day, by = RS) + RS + Date** | **9** | **-9794.03** | **0** |
| s(Day)+ Date | 6 | -6770.47 | 403.57 |
| s(Day) + RS + Date, | 7 | -6768.47 | 405.57 |
| Date | 4 | -6545.34 | 628.69. |
| RS + Date | 5 | -6543.34 | 630.69 |

**Table S3.2:** Support for the candidate generalised additive mixed models explaining variation in hourly VeDBA_activity_*.

| **Models** | **K** | **AIC** | **∆AIC** |
| --- | --- | --- | --- |
| **S(Hour, by = RS** *** period)**  **+ RS * period + Date** | **11** | 255352.0 | 0 |
| S(Hour, by = RS * period) + RS + period + Date | 10 | 255950.7 | 598.66 |
| S(Hour, by = RS * period) + RS + Date | 9 | 256294.2 | 942.13 |
| S(Hour, by = RS) **+** RS * period + Date | 9 | 256532.1 | 1180.04 |
| S(Hour, by = RS) + RS + period + Date | 8 | 257125.7 | 1773.65 |
| Date | 4 | 269619.7 | 14277.70 |

**Table S3.3:** Support for the candidate generalised additive mixed models explaining variation in the proportion of open habitat used during daytime.

| **Models** | **K** | **AIC** | **∆AIC** |
| --- | --- | --- | --- |
| **s(Day, by = RS) + RS + Date** | **8** | **69734.9** | **0** |
| s(Day) + RS + Date | 6 | 69794.8 | 59.80 |
| s(Day) + Date | 5 | 69796.4 | 61.43 |
| RS + Date | 4 | 72543.4 | 2808.42 |
| Date | 3 | 72545.6 | 2810.59 |

**Table S3.4:** Support for the candidate generalised additive mixed models explaining variation in distance to the nearest road in open habitat during daytime.

| **Models** | **K** | **AIC** | **∆AIC** |
| --- | --- | --- | --- |
| **s(Day, by = RS) + RS + Date** | **9** | **167866.0** | **0** |
| s(Day) + RS + Date | 7 | 167921.0 | 54.98 |
| s(Day) + Date | 6 | 167928.2 | 62.19 |
| RS + Date | 5 | 168005.5 | 142.48 |
| Date | 4 | 168015.6 | 149.61 |

**Table S3.5:** Support for the candidate generalised additive mixed models explaining variation in distance to the nearest woodland in open habitat during daytime.

| **Models** | **K** | **AIC** | **∆AIC** |
| --- | --- | --- | --- |
| **s(Day, by = RS) + RS + Date** | **9** | **173254.8** | **0** |
| s(Day) + Date | 6 | 173328.2 | 73.34 |
| s(Day) + RS + Date | 7 | 173329.7 | 78.91 |
| RS + Date | 5 | 173826.6 | 571.76 |
| Date | 4 | 173828.1 | 573.29 |

**Appendix S4: Model selection with AIC criteria for** **the analysis of behavioural modifications linked to female parturition date.**

Models were ranked according to AIC selection criteria using the difference in the values of AIC (∆AIC), the number of estimated parameters (K) and Akaike weights. The retained model is in bold.

**Table S4.1:** Support for the candidate linear mixed models explaining variation in daily VeDBA_activity_*.

| **Models** | **K** | **AIC** | **∆AIC** |
| --- | --- | --- | --- |
| **Date * period** | **6** | **-4529.2** | **0** |
| Date + period | 5 | -4497.8 | 31.40 |
| period | 4 | -4496.2 | 33.05 |
| Date | 4 | -4024.7 | 504.58 |
| Constant | 3 | -4023.0 | 506.23 |

**Table S4.2:** Support for the candidate linear mixed models explaining variation in the proportion of open habitat used during daytime. The most complex model included the two-way interaction between date of parturition (Date) and period, with individual identity as a random effect on the intercept.

| **Models** | **K** | **AIC** | **∆AIC** |
| --- | --- | --- | --- |
| **Date * period** | **5** | 45921.1 | 0 |
| Date + period | 4 | 46322.9 | 401.88 |
| Period | 3 | 46334.4 | 413.37 |
| Date | 3 | 47607.0 | 1685.94 |
| Constant | 2 | 47617.3 | 1696.27 |

**Table S4.3:** Support for the candidate linear mixed models explaining variation in distance to the nearest woodland in open habitat during daytime. The most complex model included the two-way interaction between date of parturition (Date) and period, with individual identity as a random effect on the intercept.

| **Models** | **K** | **AIC** | **∆AIC** |
| --- | --- | --- | --- |
| **Date * period** | **6** | 113314.8 | 0 |
| Period |  | 113337.5 | 22.80 |
| Date + period | 5 | 113338.5 | 23.74 |
| Constant | 3 | 113606.6 | 291.86 |
| Date | 4 | 113607.9 | 293.14 |

**Table S4.4:** Support for the candidate linear mixed models explaining variation in the distance to the nearest road in open habitat during daytime. The most complex model included the two-way interaction between date of parturition (Date) and period, with individual identity as a random effect on the intercept.

| **Models** | **K** | **AIC** | **∆AIC** |
| --- | --- | --- | --- |
| **Period** | **4** | **111890.6** | **0** |
| Date * period | 6 | 111892.0 | 1.38 |
| Date + period | 5 | 111892.6 | 2.00 |
| Constant | 3 | 111962.0 | 71.41 |
| Date | 4 | 111964.0 | 73.41 |
